# Suppression of Transposable Elements in Leukemic Stem Cells

**DOI:** 10.1101/169524

**Authors:** Anthony R. Colombo, Asif Zubair, Devi Thiagarajan, Sergey Nuzhdin, Timothy J. Triche, Giridharan Ramsingh

**Affiliations:** Keck School of Medicine of University of Southern California, Jane Anne Nohl Division of Hematology and Center for the Study of Blood Diseases, Los Angeles, California 90033, USA.; University of Southern California, Department of Molecular and Computational Biology, Los Angeles, CA 90089-2910, USA.; Langone Medical Center of New York University School of Medicine, Endocrinology Division for the Study of Diabetes, 550 1^st^ Avenue, New York, NY 10016, USA

## Abstract

Genomic transposable elements (TEs) comprise nearly half of the human genome. The expression of TEs is considered potentially hazardous, as it can lead to insertional mutagenesis and genomic instability. However, recent studies have revealed that TEs are involved in immune-mediated cell clearance. Hypomethylating agents can increase the expression of TEs in cancer cells, inducing ‘viral mimicry’, causing interferon signalling and cancer cell killing. To investigate the role of TEs in the pathogenesis of acute myeloid leukaemia (AML), we studied TE expression in several cell fractions of AML while tracking its development (pre-leukemic haematopoietic stem cells, leukemic stem cells [LSCs], and leukemic blasts). LSCs, which are resistant to chemotherapy and serve as reservoirs for relapse, showed significant suppression of TEs and interferon pathways. Similarly, high-risk cases of myelodysplastic syndrome (MDS) showed far greater suppression of TEs than low-risk cases. We propose TE suppression as a mechanism for immune escape in AML and MDS. Repression of TEs co-occurred with the upregulation of several genes known to modulate TE expression, such as RNA helicases and autophagy genes. Thus, we have identified potential pathways that can be targeted to activate cancer immunogenicity via TEs in AML and MDS.

## Introduction

Transposable elements (TEs) have been mostly considered detrimental because of their inherent mobile nature. Their expression can lead to insertional mutagenesis, chromosomal rearrangements, and genomic instability, potentially contributing to cancer development ^1–4^ TEs have the ability to transpose to new sites through a cut-and-paste mechanism (DNA transposons) or through RNA intermediates by a copy-and-paste mechanism (retrotransposons). Retrotransposons are further classified into long terminal repeat (LTR) and non-LTR elements. Endogenous retroviruses (ERV), which are LTRs, resemble retroviruses in their structure and function. Long interspersed nuclear elements (LINE) such as LINE1 are non-LTRs, and autonomous in their ability to retrotranspose, whereas short interspersed nuclear elements (SINE) such as *Alu* are non-autonomous, and dependent on LINE for retrotransposition. TEs are highly expressed during embryogenesis and play an active role in it^5, 6^. TEs have also been suggested to have played a positive role in evolution by increasing the potential for advantageous novel genes ^7–10^.

The genomic regions that contain TEs are highly methylated and are silenced by heterochromatin in the somatic cells^11, 12^. TE activation has been reported in aging tissues, including in aging stem cells^13, 14^. TEs have been reported to be expressed in various types of cancers for the past 3 decades; however, it remains unknown if they are causal or consequential to the development of cancer. Recent reports revealed a potential beneficial role of TEs in cancer, wherein ERVs were shown to be potential tumour-specific antigens^15^. Hypomethylating agents increase the expression of TEs in cancer cells, inducing ‘viral mimicry’ and causing interferon signalling and cancer cell killing^16, 17^ Bidirectional (sense and anti-sense) transcription of many TEs, including ERVs, yields dsRNA^18, 19^. dsRNA sensors then activate potent interferon response pathways, leading to the activation of inflammatory pathways and cell death^16, 17^ These findings suggested that TE expression in cancer cells could play a role in immune-mediated clearance of cancer cells.

Acute myeloid leukaemia (AML), the most common form of acute leukaemia in adults, is characterized by high rates of initial remission with chemotherapy (60-70%), but is also associated with high relapse rates. Nearly two decades ago, it was shown that only a small fraction of AML cells (termed leukemic stem cells or LSCs) were capable of re-initiating the tumour when transplanted into immunodeficient animals^20^. LSCs in AML can be identified based on the expression of cell surface proteins (CD34^+^CD38^neg^CD99^+^TIM3^+^)^21^. Although the exact role of LSCs in the pathogenesis and relapse of AML is still debated, their presence is associated with resistance to therapy, relapse, and poor prognosis^22^. Thus targeting LSCs in AML is a major focus of oncologic research, however the lack of understanding of pathways dysregulated in LSCs has hampered progress. We speculated that the resilience of LSCs was mediated by its ability to escape immune mediated clearance. To investigate this, we studied the expression of TEs and its accompanying immune pathways in AML cell fractions.

## Materials and methods

See supplemental section for materials and methods.

## Results

### LSCs show low expression of TEs

Corces et al. had recently used fluorescent activated cell sorting to isolate leukemic cells from patients with AML. They separated the cells of three distinct stages of AML evolution, pre-leukemic haematopoietic stem cells (pHSCs; CD34^+^CD38^neg^CD99^neg^TIM3^neg^), leukemic stem cell (LSCs; CD34^+^CD38^neg^CD99^+^TIM3^+^), and leukemic blasts (Blasts; CD99^+^TIM3^+^CD45^mid^SSC^high^), characterized their transcriptome, and analysed their coding gene expression patterns^21^. To investigate the regulation of TEs in the development of AML, we examined the transcriptomes in these stages by measuring the changes in TE expression. When LSCs were compared to pHSCs and Blasts, we identified a significant downregulation of TEs in LSCs (Figure 1A, Figure 1B, and Supplement Figure 1). Among the different classes of TEs, SINE was the most suppressed in LSCs, followed by LTR retrotransposons (Figure 1A). The most dysregulated TE types in LSCs were *Alu*, ERV1, ERVL, ERVK, and LTR retrotransposons, all of which showed significant suppression (Table 1).

**Figure 1:**
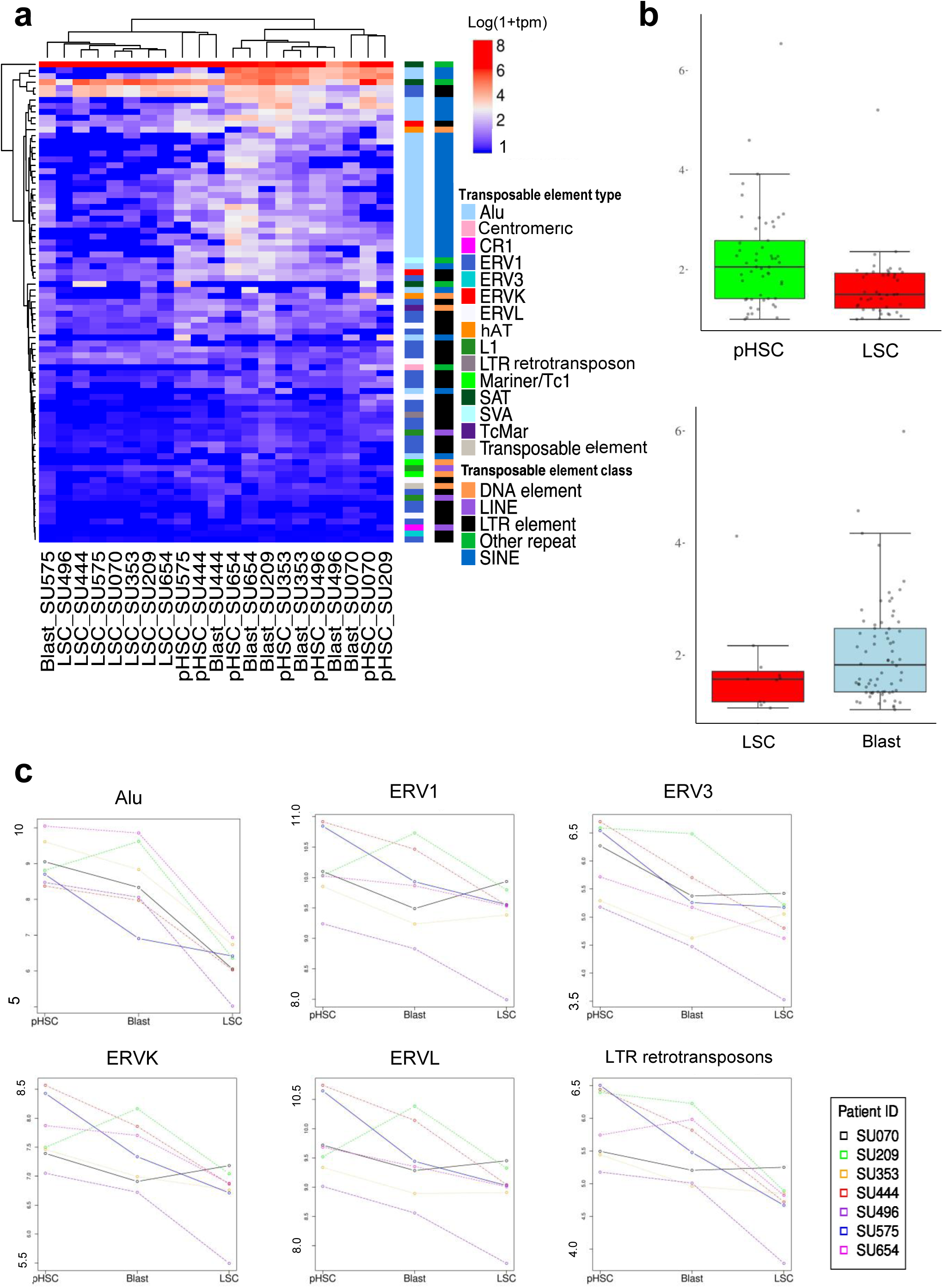
Analysis of differential expression of transposable elements in pre-leukemic stem cells (pHSC), leukemic stem cells (LSC), and Blasts. A) X-axis; Patient identifier. The expression levels in log_10_ using the metric transcripts per million (TPM). The ‘Transposable Element (TE) Type’ classifies individual repeat transcripts into one of 68 unique canonical categories of TEs. Each TE type is contained in one TE Class. B) Quantiles of the absolute log-fold change of the differentially expressed (DE) TE transcripts in pHSC-LSC, and Blast-LSC samples. Y-axis: absolute log-fold change of each individual DE TE transcript from Figure 1A. C) Y-axis: log_10_ of TPM expression level for each of the 7 paired samples across each clonal point. The individual patients are denoted with unique colours.

We further analysed the dysregulation of TEs in individual AML samples, while tracking the stages of AML. We found that specific TE types were dysregulated, with LSCs showing significant suppression of *Alu*, ERV3. ERVK, ERVL, and LTR retrotransposons (Figure 1C, Supplement Figure 2). We did not observe significant suppression of LINE1 in LSCs. These results suggested that TEs were dysregulated during AML development, with LSCs showing significant suppression of specific TE types.

### LSCs show suppression of interferon pathways

LSCs are known to be resistant to treatment and serve as potential sources of relapse for AML, although the mechanisms behind this resilience are not fully understood^22^. Expression of TEs is known to activate a viral recognition pathway, which causes interferon signalling and immune-mediated cell clearance^16, 17^. Because LSCs showed suppressed TE expression, we investigated whether this TE suppression was associated with the suppression of interferon pathways in LSCs, which could enable its escape from immune-mediated clearance. LSCs showed significantly higher suppression of several Gene Ontology Consortium (GO)-interferon signalling pathways than Blasts (Figure 2A). When immune-related pathways (with a set of 335 genes, generated by combining 17 canonical immune pathways in MSigDB) and inflammatory pathways (with a set of 649 genes combining acute inflammatory response and inflammatory response in MSigDB and GO) in LSCs and Blasts were compared, LSCs showed significant suppression of the immune-related pathways (Figure 2B, Supplement Figure 3, and Supplement Table 5).

**Figure 2:**
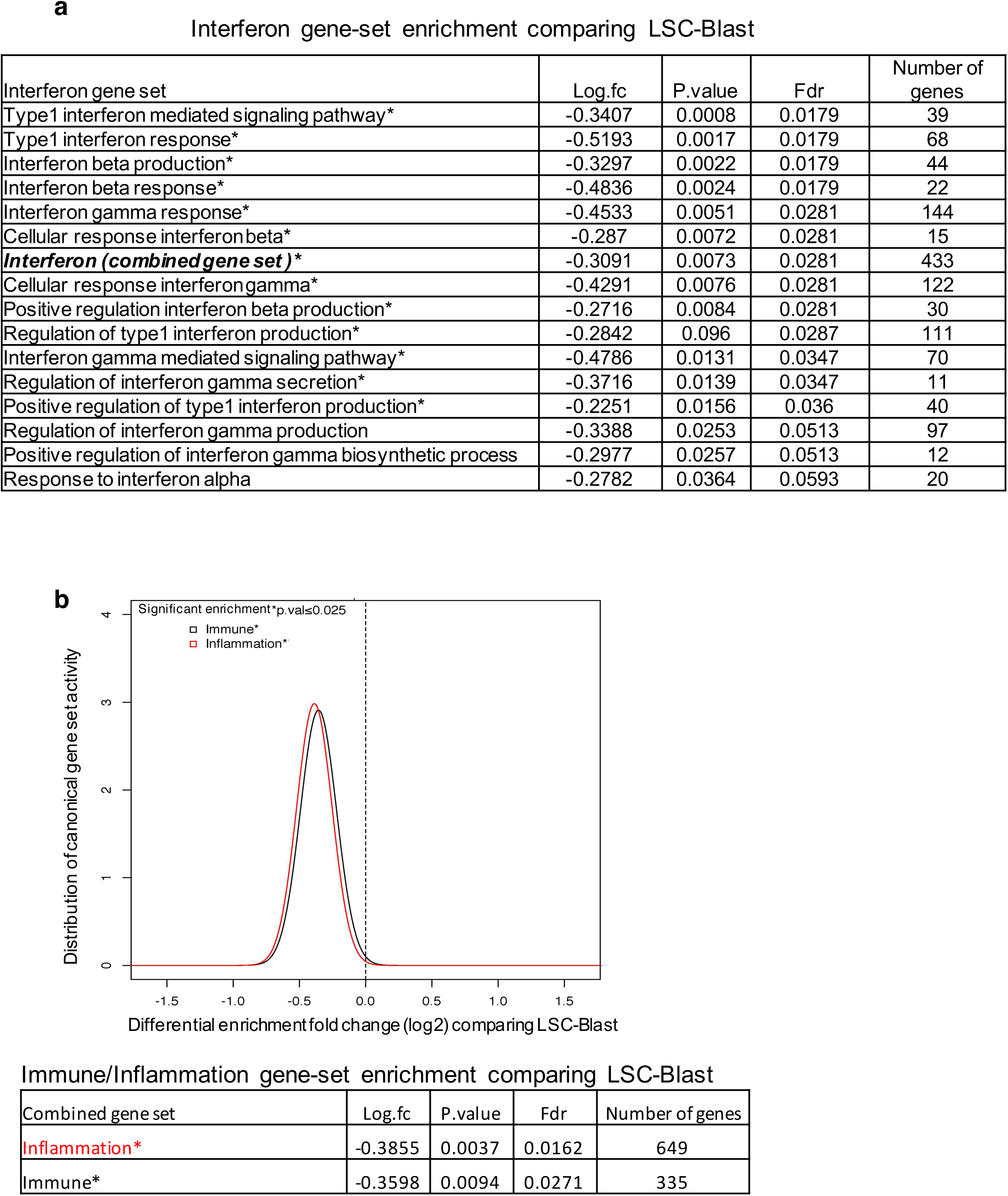
Analysis of gene set enrichment for interferon, inflammation, and immune response genes. A) Interferon-related gene sets from GO MSigDB, comparing LSCs and Blasts in AML. B). Gene set enrichment analysis of combined inflammation and immune gene sets, comparing LSCs and Blasts. The Bonferroni multiple testing correction significance threshold is denoted as ‘p.val’. ^∗^ indicates p < 0.025.

However, comparison between LSCs (which showed lower expression of TEs than pHSCs) and pHSCs showed no significant differences in interferon, immune, or inflammatory pathways (Supplement Figure 3). This appeared to contradict the model of TE-induced activation of immune pathways. We therefore investigated alternate pathways that could suppress in immune pathways in pHSCs. Interestingly, we found that all pHSCs exhibited very high expression of EVI-1 (pHSCs vs. LSCs, 4.6-fold, p < 0.0001; pHSCs vs. Blasts, 3.4-fold, p < 0.0001, Supplement Figure 4), which is known to suppress immune pathways by downregulating NFκB (a pathway known to be activated by viral RNA) ^23^. Consistent with this finding, we also observed that NFκB pathways were more suppressed in pHSCs than Blasts (Blasts and pHSCs showed similar expression of TEs) (Supplemental Figure 4). These findings suggested that both LSCs and pHSCs showed suppression of NFκB and immune-related pathways, compared to Blasts. LSCs showed suppressed TE expression and pHSCs showed high expression of EVI-1.

### Coding gene networks are co-regulated with TEs

Although TE expression is known to activate immune pathways, the types of TEs that participate in this mechanism are currently unknown. In order to understand the relationship between coding gene expression and the expression of specific TE subtypes, we first performed an unsupervised clustering of the AML samples based on coding gene expression, and found that LSCs formed a well-grouped cluster (Figure 3). We then analysed the corresponding expression of various TE types and observed a significant suppression in the expression of specific TE types such as *Alu*, ERV3, ERVK, and LTR retrotransposons in LSCs, compared to pHSCs and Blasts (Figure 3). This suggested that coding gene expression was distinct in samples with low expression of specific types of TEs (LSCs).

**Figure 3:**
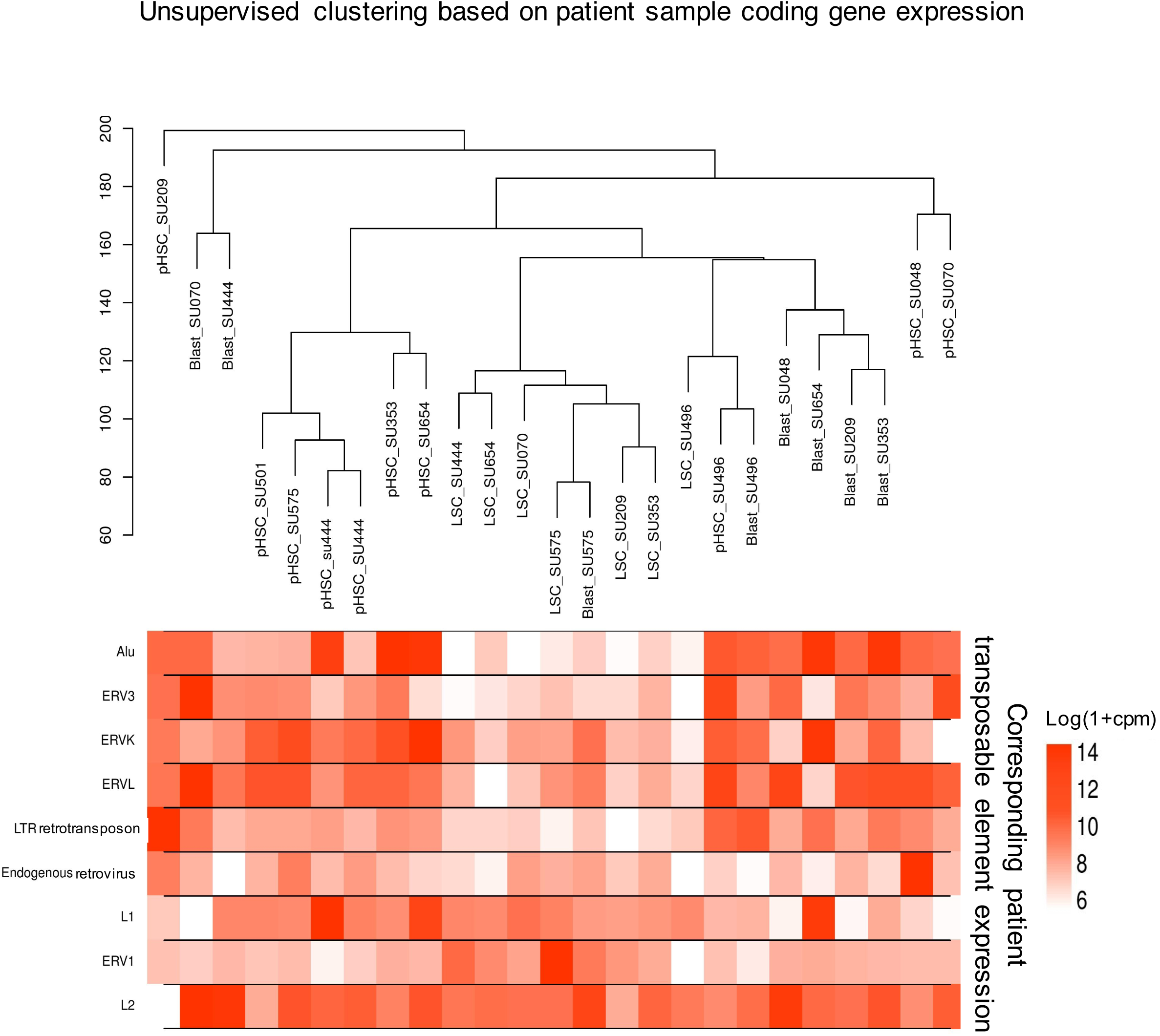
Unsupervised hierarchical clustering of coding gene expression in patient samples and the expression levels of the corresponding transposable elements. A) The image on top depicts the hierarchical clustering of each group (pHSC, LSC, and Blast) based on the average Euclidean distance for the coding gene expression in the patient samples. Below each sample, the expression of the corresponding TE Types (Alu, ERV3, ERVK, ERVL, LTR Retrotransposon, Endogenuous Retroviruses, L1, ERV1, and L2) is shown. The TE expression is expressed in units of normalized counts per million (CPM) of log_10_ (1 + CPM).

Next, in order to investigate which coding gene networks were correlated with specific TE types, we created a genomic association table using the transcriptome from Blasts and LSCs, as shown in Figure 4. The coding genes were first clustered based on their co-expression to form specific modules. Each module contained unique set of genes that were likely co-regulated and had functional similarities. For example, module 26 contains many RNA helicase genes (Supplement Figure 5). We correlated these modules to the expression of specific TE types and found that some modules were positively or negatively correlated with the expression of specific TE types. We performed a pathway analysis using the genes in each module for testing the interferon, immune and inflammatory activity, comparing Blasts to LSCs. We identified modules that showed activation (modules 3, 5, 13, 14, 17 and 41) and suppression (22, 24, 26, 29, 30, 39 and 46) of interferon/immune/inflammation gene pathways in Blasts, compared to LSCs (Figure 4 and Supplement figure 5). We then correlated this with the expression of different TE types. As shown in the Figure 4, the modules that had shown activation of interferon/immune/inflammation genes were positively associated with the expression of specific TE types (*Alu*, ERVL, ERVK, and LTR retrotransposons) and negatively associated with the expression of ERV1, SAT, and L1. The modules that had shown suppression of the genes in interferon/immune/inflammation were positively associated with the expression of ERV1 and negatively associated with *Alu*, ERV3, ERVL, and LTR retrotransposons. Chi-square test confirmed a global association between the correlation of positive/negative coding gene module with TE types and the positive/negative enrichment activity of the interferon/immune/inflammation pathways, respectively (p =0.005). This suggested that specific types of TE were significantly linked to interferon/immune/inflammatory pathway activation.

**Figure 4:**
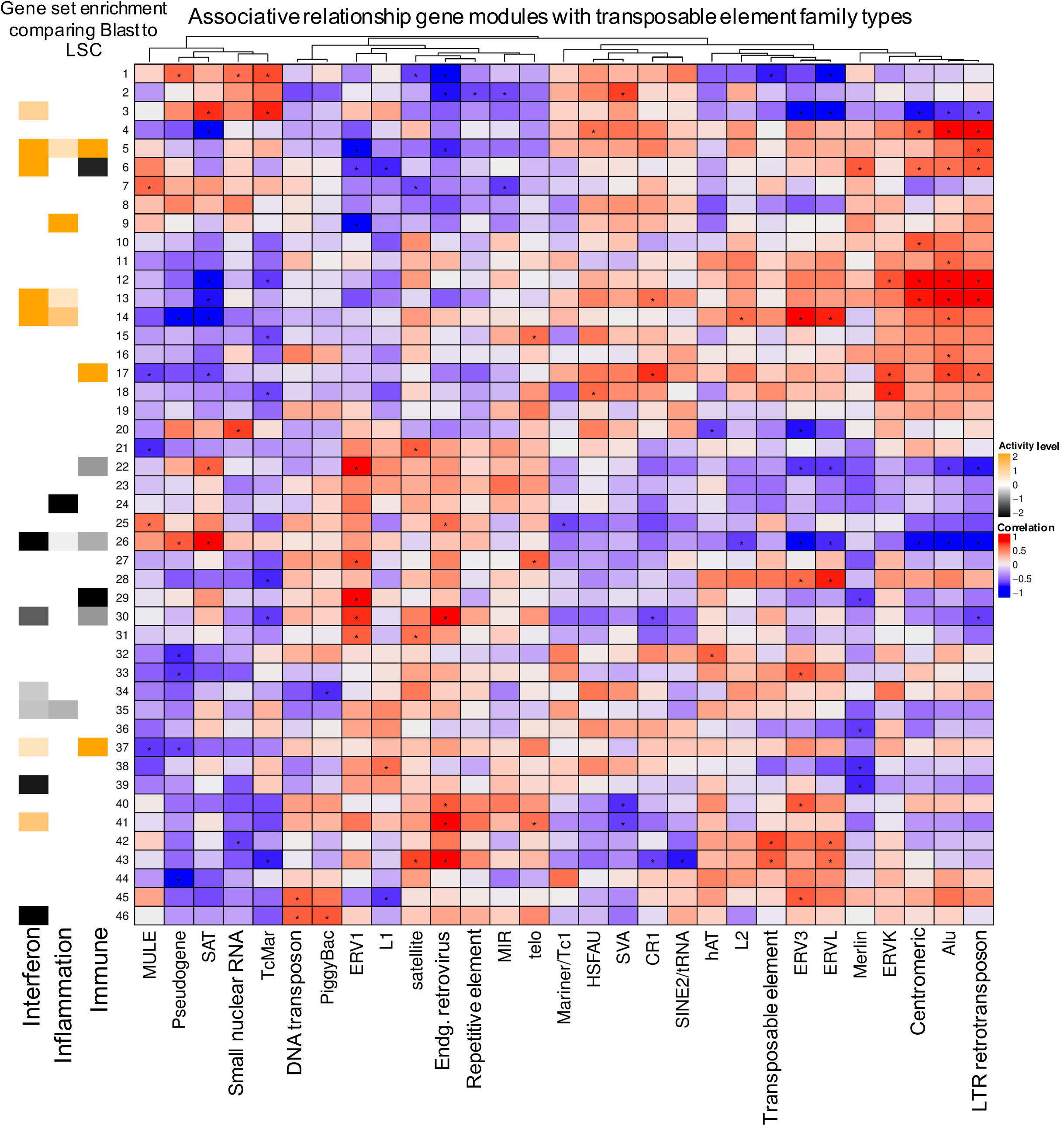
Identifying significant associations between the expression of coding gene network and the transposable element types in AML. The numbers on the Y-axis denote the gene ‘modules’ constructed by identifying gene networks base on co-expression patterns. The X-axis denotes canonical TE types used for correlating them. The centre figure of squares represents the correlation matrix for the normalized gene ‘module’ expressic and the TE type. ^∗^ indicates significant associations (p.value ≤ 0.05). Each gene ‘module’ was tested for activation of canonical immune and inflammation gene sets in Blasts and compared with LSCs. The significant (p.value ≤ 0.05) pathway activity level for each module is plotted on the left of Y-axis (yellow indicates significantly higher activation in Blast, and black indicates significantly higher activity in LSCs).

### High-risk cases of MDS show suppression of TEs

In order to validate the observation that TEs induced inflammatory pathway activation in Blasts in an independent model, we analysed the expression of TEs in MDS, comparing CD34+ cells from low-risk and high-risk cases of MDS. MDS cases with refractory anaemia with excess blasts (RAEB) were classified as high-risk and the others were considered low-risk. The two groups were compared using RNA sequencing data from Wang et al^24^. We identified significant suppression of TE expression in high-risk MDS, compared to low risk MDS (Figure 5A). High-risk MDS specifically showed suppression of Type 1 interferon genes, which are known to be activated by viral RNA (Figure 5B). Inflammation-related genes (Figure 5C, 2-fold change, p = 0.0002, FDR = 0.0003 and Supplement figure 6) were also significantly suppressed in high-risk MDS, compared to low risk-MDS. The model validated many of the features of AML development (Figure 1, Figure 2), where suppression of TEs is associated with diminished expression of interferon and inflammatory genes.

**Figure 5:**
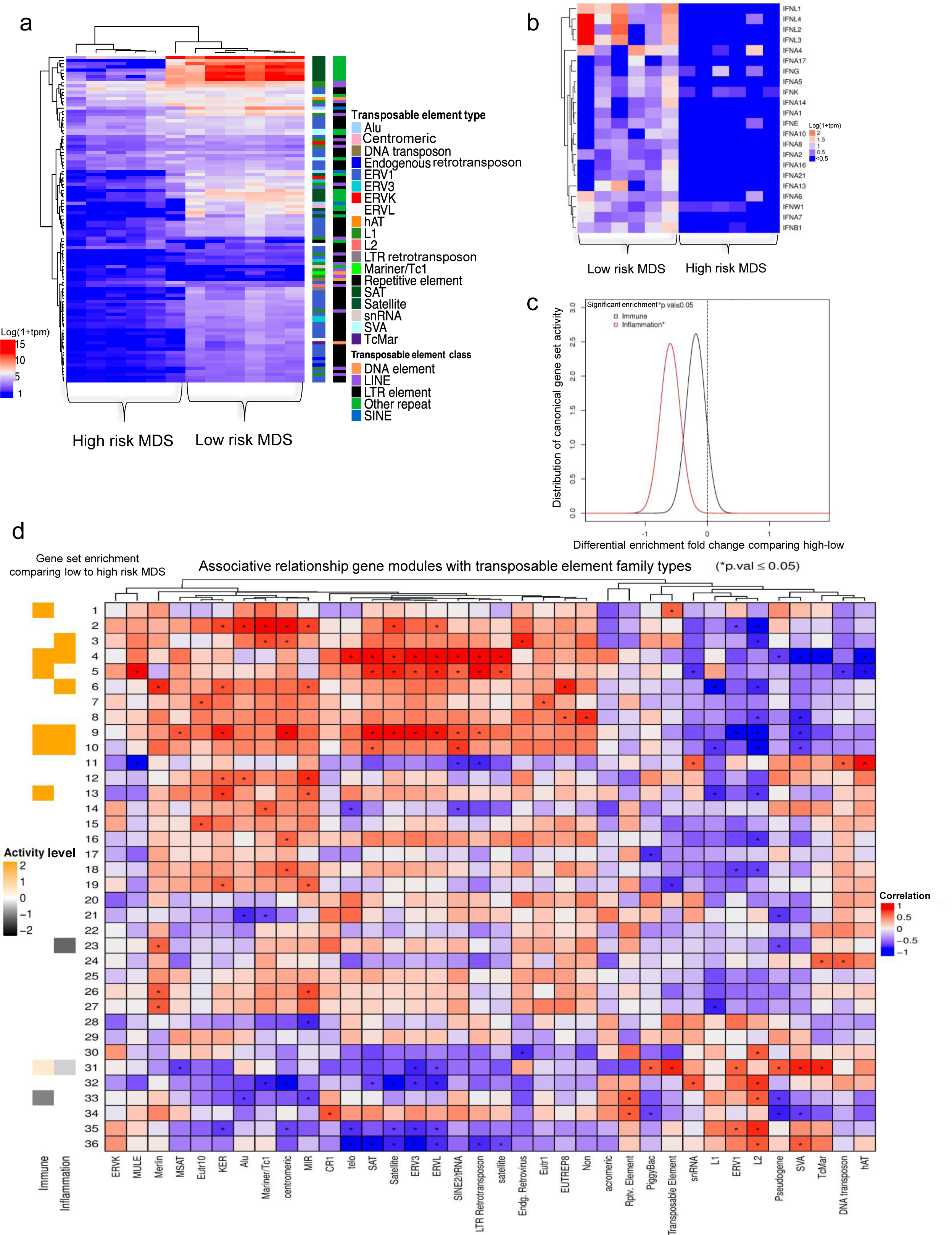
Expression of transposable elements in myelodysplastic syndrome (MDS) A) Differential TE expression between low- and high-risk MDS cases. The legends showing TE type and class are identical to Figure 1A. B) Expression of type-1 interferon genes in MDS using log_10_ TPM C) Comparison of high- and low-risk MDS cases for enrichment of canonical immune and inflammation gene sets, similar to Figure 2B. D) Identification of significant association between the expression of coding gene networks and TE types in MDS, similar to Figure 4.

To further characterise the association between coding genes and TE expression, we created an association table similar to that for AML shown in Figure 4 (Figure 5D). The data indicated a similar association between gene modules (module 3, 4, 5, 9 and 10) that showed activation of immune/inflammatory genes in low-risk MDS compared to high-risk MDS and the expression of specific TE types such as ERV3, ERVL, and LTR retrotransposons. These modules also showed a negative correlation with the expression of ERV1 and L1 (Figure 5D and Supplement Figure 7). Type 1 interferon genes were present in only module 9 and 10. These data indicated that similar to LSCs in AML, high-risk cases of MDS exhibited suppressed expression of specific TE types along with the corresponding suppression of interferon and inflammatory pathways.

### Pathways that potentially mediate suppression of TEs in LSCs

The mechanisms behind the regulation of TEs have not been thoroughly investigated. Similar to coding genes, TEs can be regulated both transcriptionally and post-transcriptionally. Epigenetic modifications secondary to alterations in ATRX, P53, and SIRT1 and methylation of DNA, have been shown to regulate the expression of TEs^25–27^. We investigated whether TEs were suppressed in LSCs through epigenetic mechanisms by analysing its chromatin accessibility using the data from assay for transposase accessible chromatin with high-throughput sequencing (ATAC-seq) for pHSCs, LSCs, and Blasts from Corces et al^21^. ATAC-seq has been used for genome-wide mapping of chromatin accessibility. It uses Tn5 transposase to insert sequencing adapters into accessible regions of the chromatin and then uses the sequence reads mapped to the genome to infer accessible regions. Principle component analysis showed that pHSCs were clustered separately from LSCs and Blasts (Figure 6A). Contrary to our expectations, LSCs, despite having low expression of TEs, had more nucleosome-free regions than pHSCs (Figure 6B). We analysed the differential accessibility by comparing the accessibility of LSCs to pHSCs, and found 18,099 regions that were significantly more accessible and 441 regions that were significantly less accessible in LSCs compared to pHSCs (Figure 6B and Supplement Table 3). Comparison of LSCs to Blasts showed no significant differences in the accessible regions. These findings suggested that the suppression of TEs in LSCs was likely not due to increased heterochromatin.

**Figure 6:**
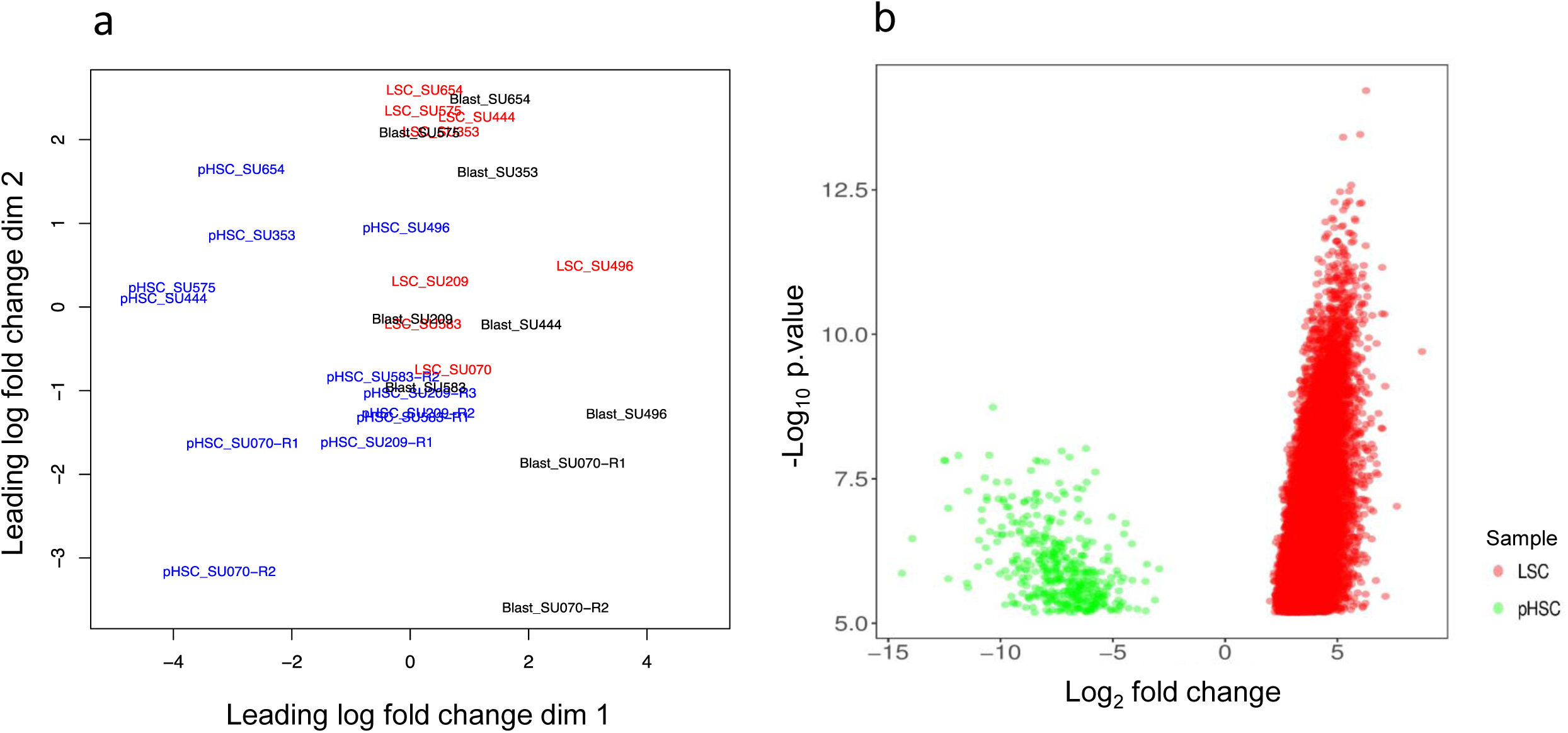
Chromatin accessibility in pHSCs, LSCs, and Blasts. A) Multi-dimensional scaling plot with two dimensions showing similarity between different ATACseq samples: pHSC (blue), LSC (red), and Blast (black). B) Depicts the differential accessibility using ATACseq sampling data comparing LSCs to pHSC. X-axis is log_2_ fold change of differentially accessible regions (Supplement table 4); Y-axis is –log_10_ of the p.values reported from comparison. The minimum p.value considered was 5.593e-03.

Because LSCs showed suppressed TE expression despite having more accessible chromatin, we investigated other pathways that could regulate TE expression. A major mechanism for regulating TEs involves their post-transcriptional degradation^28–30^. We analysed genes known to suppress TEs post-transcriptionally, as described by Goodier et al. ^28^, and compared them in LSCs and Blasts and in high-risk and low-risk MDS. High-risk MDS showed significant upregulation of ATG5, KIAA0430, CALCOCO2, ZC3HAV1, HNRNPL, and PABPC1, compared to low-risk cases. LSCs showed significant upregulation of ATG5 and KIAA0430 (Figure 7A, Supplement Figure 8). High-risk MDS cases also showed significant upregulation of RNA interference genes such as DROSHA, DICER1, and DGCR8, compared to the low-risk cases, but they were not significantly upregulated in LSCs (Figure 7A). Similar to the piRNA system in males, KIAA0430 or meiosis arrest female protein 1 is known to play a key role in repressing TEs during oogenesis^31^. However, its role in regulating TEs in somatic cells has not been reported. Autophagy-related 5 (ATG5), which was significantly upregulated in both LSCs and high-risk MDS cases (Supplement figure 8), mediated autophagy by enabling the formation of autophagy vesicles. Autophagy is a process by which various intracellular components are transported to the lysosomes and degraded. A recent study showed that autophagy mediates the degradation of TE post-transcriptionally^32^. Interestingly, LAMP2 was also upregulated in both LSCs and high-risk MDS cases (Figure 7B). Recently, it was shown that LAMP2C, a splice isoform of LAMP2, mediated the degradation of RNA via autophagy (RNAutophagy) ^33–35^. HSP90AA1 (heat shock protein 90 kDa α [cytosolic], class A member 1) is a pathogen receptor that activates autophagy and thus controls the viral infection^36^. This protein was also seen upregulated in high-risk MDS cases and LSCs (Figure 7A and B).

**Figure 7:**
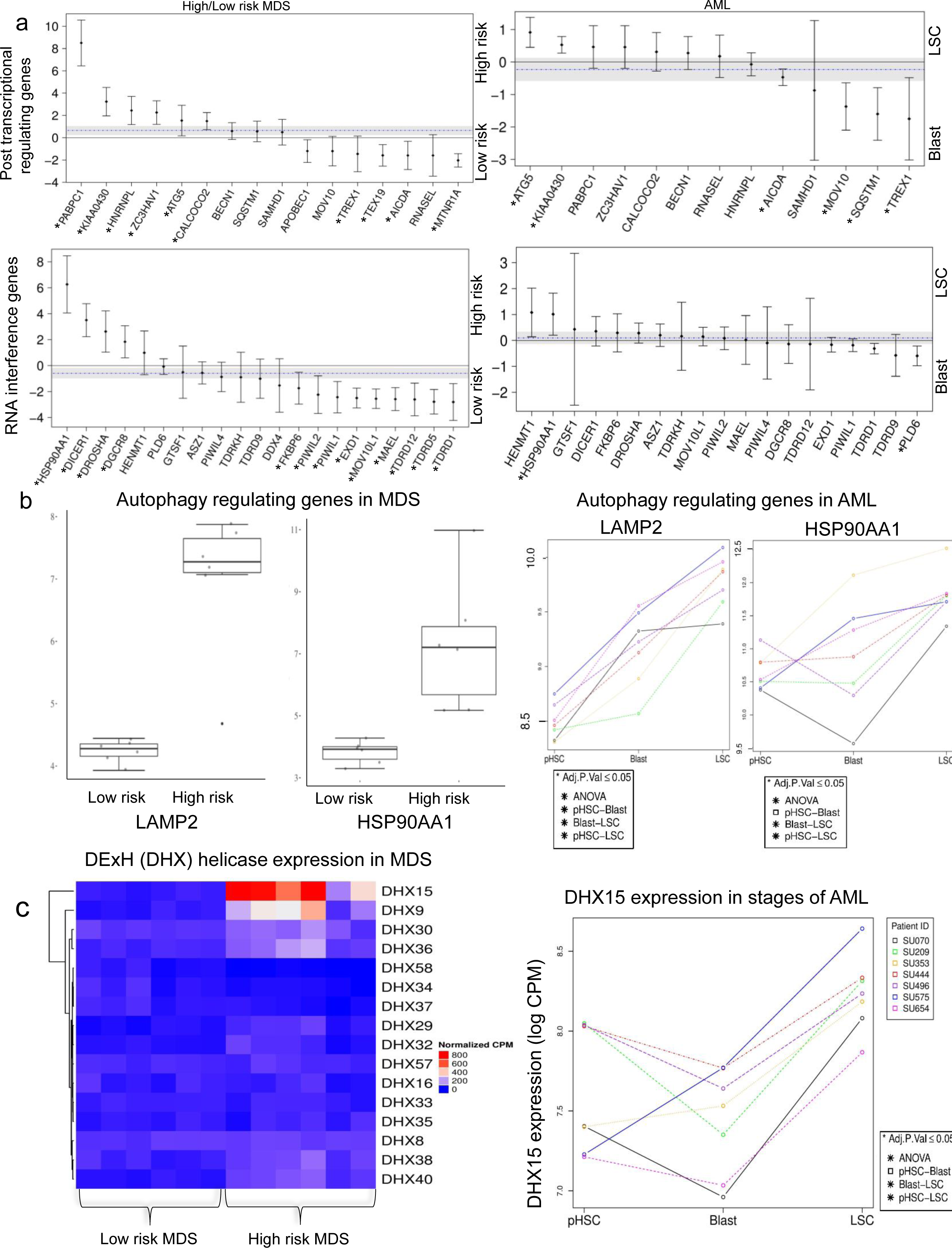
Expression of genes that modulate transposable elements post-transcriptionally. A) Genes that regulate TE post-transcriptionally. Positive fold-change (Y-axis; y > 0) indicates higher expression in high-risk MDS cases and/or LSCs. Negative fold-change (y < 0) indicates higher expression in low-risk MDS cases and/or Blasts. Significant genes are denoted with ^∗^p.value ≤0.05. B) Autophagy-regulating genes in MDS and AML. Expression of LAMP2 and HSP90AA1 in high- and low-risk MDS cases and pairwise comparison of AML stages. Paired patient measurements are shown with matching colours. Adjusted significance values denoted ^∗^p.value ≤0.05 C) Expression of RNA helicase genes, DExH genes. The heatmap depicts differentially expressed DExH genes in MDS cases. Expression of DHX15 in different stages of AML, where paired patient measurements are shown with matching colours. Adjusted significance values denoted ^∗^p.value ≤0.05

RNA helicases are known to bind to and degrade TE post-transcriptionally^28, 29, 37–40^. We found significant upregulation of the DExH class of RNA helicases (DHX) in high-risk MDS cases (Figure 7C and Supplement figure 8). In particular, DHX15 and DHX9 were almost exclusively expressed in high-risk MDS cases and DHX15 was significantly upregulated in LSCs, compared to Blast (Figure 7C and Supplement figure 8). These results indicated the possibility that several post-transcriptional mechanisms operated for mediating the suppression of TEs in AML and MDS.

### Discussion

Our study is the first to comprehensively evaluate the expression of TEs and its association with coding genes in cancer. We demonstrated that the expression of TEs was dysregulated during the development of AML and MDS, with LSCs and high-risk MDS showing significant suppression. It has been shown that the suppression of the viral recognition pathway conferred resistance to chemotherapy; mutations in MAVS and RIG-1, genes in the viral recognition pathway, have been reported in cancer^41^. We speculated that the expression of TEs could be a potential mechanism for immune-mediated elimination of cancer cells.

Hypomethylating agents have been found to be useful for treating AML and MDS, and recent studies have reported that the activation of TEs with the subsequent immune activation was important for their efficacy against cancers^16, 17^. Here, we demonstrated that these mechanisms likely operated naturally during cancer development and progression to enable immune-mediated control of AML and MDS. Despite the efficacy of hypomethylating agents against AML and MDS, only a minority (~20%) of patients responded to this therapy^42^. Among patients who did respond, most eventually developed resistance to therapy with hypomethylating agents. Understanding the regulation of TEs would help us explore predictive factors for hypomethylating treatment and develop novel strategies to prevent relapse in patients treated with hypomethylating agents.

The role of LSCs in the pathogenesis of AML remains controversial. Our results showed that LSCs clearly suppressed the expression of TEs along with distinct coding gene expression. They also showed more suppression of inflammatory pathways, including the NFκB pathway. Since Blasts are short lived, they probably did not evolve mechanisms to escape immune-mediated attacks. We speculate that LSCs are a subset of Blasts with the ability to evade immune recognition.

pHSCs, despite having similar expression levels of TEs as Blasts, also showed suppression of inflammatory pathways that prevent the activation of immune signalling. EVI-1, which is known to suppress NFκB, was uniquely over-expressed in pHSCs, suggesting that there exists distinct genes which suppress the inflammatory pathways in pHSCs. pHSCs carry mutations in genes regulating the epigenetic machinery and have been clearly demonstrated to precede the development of AML^43^. pHSCs are resistant to chemotherapy and likely function as reservoirs for relapse of leukaemia^44, 45^. High expression of TEs in pHSCs makes them vulnerable to clearance through the viral-recognition pathway; however, this event likely never occurs because of EVI1-mediated suppression of NFkB, which is downstream to the viral-recognition pathway. High expression of EVI-1 has been shown to be an indicator of poor risk in AML^46^. Our analysis is the first to highlight that EVI-1 was significantly expressed at high levels in pHSCs. Targeting EVI-1 in pHSCs could help prevent clonal evolution in AML. For example, miR-133 is known to target EVI-1^47^. It would be important to explore its role in clonal haematopoiesis in the elderly, a condition characterized by expansion of haematopoietic stem cells with mutations in pHSCs.

Our analysis that correlated the expression of coding gene networks to the expression of TE types revealed an association between inflammatory pathways to SINE and LTR families and an anti-association with LINE1. Among the types of TEs, LINE1 is known to have the highest activity of retrotranspositioning and thus has the most potential to cause genomic instability. Hence, LSCs might have co-opted to evolve by suppressing the inflammation-inducing TE classes, while retaining the expression of LINE1, which could potentiate genomic instability and hence clonal evolution.

We found high expression of several DExH RNA helicases in high-risk MDS, but their role in regulating TEs has not yet been reported. RNA helicases bind to single as well as double stranded RNA, and regulate gene splicing. Aberrant splicing events have been reported in patients with MDS, but it is not known whether these splicing factors also regulate TEs. Exploring this function of RNA helicases would enable us to develop drugs targeting them to activate TEs in AML and MDS.

The role of autophagy in protecting cancer cells from immune attacks via suppressing TE needs to be explored. Drugs targeting autophagy, RNA autophagy (mediated by LAMP2C) in particular, could be promising therapeutic agents against AML and MDS.

Immuno-oncology is emerging as one of the cornerstones of treatment of various cancers. Interferons have long been used in the treatment of cancers, leading to sustained remissions^48–50^. However, it has been associated with significant systemic toxicities. Activating suppressed TEs, which are known to activate interferons, in cancer cells could potentially accomplish this in a targeted manner.

Our study is the first to show dysregulation of TE in LSCs, revealing its importance in the pathogenesis of AML and MDS. Studying direct mechanisms of the regulation of cancer immunosurveillance by TEs in AML and MDS could lead to therapies improving long-term survival by manipulating the expression of TEs in leukemic cells.

## Acknowledgements

This project was funded by grants from the Leukemia Lymphoma Society-Quest for Cures #0863-15, STOP Cancer, Tower Cancer Research Foundation, and the Keck School of Medicine of the University of Southern California, Jane Anne Nohl Division of Hematology and Center for the Study of Blood Diseases, University of Southern California.

## Author Contributions

AC wrote the manuscript, analysed the experiments, and developed analytical bioinformatics software and revised the biological model. AZ developed the ATACseq software, processed, and analysed the ATACseq experimental data. DT revised the theoretical model in regard to the regulation of TE elements including key genes relating to autophagy. SN provided insight furthering the theoretical framework with regard to the role of helicases in mediating TE. TJ developed bioinformatics software. GR developed the theoretical model, analysed the experiments and wrote the manuscript.

## References

1. Belancio, V. P. LINE-1 activity as molecular basis for genomic instability associated with light exposure at night. Mob Genet. Elements 5, 1–5 (2015).

2. Kemp, J. R. & Longworth, M. S. Crossing the LINE Toward Genomic Instability: LINE-1 Retrotransposition in Cancer. Front. Chem. 3, 68 (2015).

3. Mills, R. E., Bennett, E. A., Iskow, R. C. & Devine, S. E. Which transposable elements are active in the human genome? Trends Genet. 23, 183–191 (2007).

4. Luzhna, L., Ilnytskyy, Y. & Kovalchuk, O. Mobilization of LINE-1 in irradiated mammary gland tissue may potentially contribute to low dose radiation-induced genomic instability. Genes Cancer. 6, 71–81 (2015).

5. Gerdes, P., Richardson, S. R., Mager, D. L. & Faulkner, G. J. Transposable elements in the mammalian embryo: pioneers surviving through stealth and service. Genome Biol. 17, 100-016-0965-5 (2016).

6. Ge, S. X. Exploratory bioinformatics investigation reveals importance of “junk” DNA in early embryo development. BMC Genomics 18, 200-017-3566-0 (2017).

7. Elbarbary, R. A., Lucas, B. A. & Maquat, L. E. Retrotransposons as regulators of gene expression. Science 351, aac7247 (2016).

8. Sundaram, V. et al. Widespread contribution of transposable elements to the innovation of gene regulatory networks. Genome Res. 24, 1963–1976 (2014).

9. Thompson, P. J., Macfarlan, T. S. & Lorincz, M. C. Long Terminal Repeats: From Parasitic Elements to Building Blocks of the Transcriptional Regulatory Repertoire. Mol. Cell 62, 766–776 (2016).

10. Lee, H. E., Ayarpadikannan, S. & Kim, H. S. Role of transposable elements in genomic rearrangement, evolution, gene regulation and epigenetics in primates. Genes Genet. Syst. 90, 245–257 (2015).

11. Schulz, W. A., Steinhoff, C. & Florl, A. R. Methylation of endogenous human retroelements in health and disease. Curr. Top. Microbiol. Immunol. 310, 211–250 (2006).

12. Groh, S. & Schotta, G. Silencing of endogenous retroviruses by heterochromatin. Cell Mol. Life Sci. (2017).

13. Wang, J. et al. Inhibition of activated pericentromeric SINE/Alu repeat transcription in senescent human adult stem cells reinstates self-renewal. Cell. Cycle 10, 3016–3030 (2011).

14. Sun, D. et al. Epigenomic profiling of young and aged HSCs reveals concerted changes during aging that reinforce self-renewal. Cell. Stem Cell. 14, 673–688 (2014).

15. Mullins, C. S. & Linnebacher, M. Endogenous retrovirus sequences as a novel class of tumor-specific antigens: an example of HERV-H env encoding strong CTL epitopes. Cancer Immunol. Immunother. 61, 1093–1100 (2012).

16. Chiappinelli, K. B. et al. Inhibiting DNA Methylation Causes an Interferon Response in Cancer via dsRNA Including Endogenous Retroviruses. Cell 162, 974–986 (2015).

17. Roulois, D. et al. DNA-Demethylating Agents Target Colorectal Cancer Cells by Inducing Viral Mimicry by Endogenous Transcripts. Cell 162, 961–973 (2015).

18. Lehner, B., Williams, G., Campbell, R. D. & Sanderson, C. M. Antisense transcripts in the human genome. Trends Genet. 18, 63–65 (2002).

19. Yelin, R. et al. Widespread occurrence of antisense transcription in the human genome. Nat. Biotechnol. 21, 379–386 (2003).

20. Bonnet, D. & Dick, J. E. Human acute myeloid leukemia is organized as a hierarchy that originates from a primitive hematopoietic cell. Nat. Med. 3, 730–737 (1997).

21. Corces, M. R. et al. Lineage-specific and single-cell chromatin accessibility charts human hematopoiesis and leukemia evolution. Nat. Genet. 48, 1193–1203 (2016).

22. Reinisch, A., Chan, S. M., Thomas, D. & Majeti, R. Biology and Clinical Relevance of Acute Myeloid Leukemia Stem Cells. Semin. Hematol. 52, 150–164 (2015).

23. Xu, X. et al. EVI1 acts as an inducible negative-feedback regulator of NF-kappaB by inhibiting p65 acetylation. J. Immunol. 188, 6371–6380 (2012).

24. Wang, H., Wen, J., Chang, C. C. & Zhou, X. Discovering transcription and splicing networks in myelodysplastic syndromes. PLoS One 8, e79118 (2013).

25. Leonova, K. I. et al. p53 cooperates with DNA methylation and a suicidal interferon response to maintain epigenetic silencing of repeats and noncoding RNAs. Proc. Natl. Acad. Sci. U. S. A. 110, E89–98 (2013).

26. Elsasser, S. J., Noh, K. M., Diaz, N., Allis, C. D. & Banaszynski, L. A. Histone H3.3 is required for endogenous retroviral element silencing in embryonic stem cells. Nature 522, 240–244 (2015).

27. Van Meter, M. et al. SIRT6 represses LINE1 retrotransposons by ribosylating KAP1 but this repression fails with stress and age. Nat. Commun. 5, 5011 (2014).

28. Goodier, J. L. Restricting retrotransposons: a review. Mob DNA 7, 16-016-0070-z. eCollection 2016 (2016).

29. Goodier, J. L., Cheung, L. E. & Kazazian, H. H.,Jr. MOV10 RNA helicase is a potent inhibitor of retrotransposition in cells. PLoS Genet. 8, e1002941 (2012).

30. Goodier, J. L., Cheung, L. E. & Kazazian, H. H.,Jr. Mapping the LINE1 ORF1 protein interactome reveals associated inhibitors of human retrotransposition. Nucleic Acids Res. 41, 7401–7419 (2013).

31. Su, Y. Q., Sun, F., Handel, M. A., Schimenti, J. C. & Eppig, J. J. Meiosis arrest female 1 (MARF1) has nuage-like function in mammalian oocytes. Proc. Natl. Acad. Sci. U. S. A. 109, 18653–18660 (2012).

32. Guo, H. et al. Autophagy supports genomic stability by degrading retrotransposon RNA. Nat. Commun. 5, 5276 (2014).

33. Fujiwara, Y. et al. Discovery of a novel type of autophagy targeting RNA. Autophagy 9, 403–409 (2013).

34. Fujiwara, Y., Hase, K., Wada, K. & Kabuta, T. An RNautophagy/DNautophagy receptor, LAMP2C, possesses an arginine-rich motif that mediates RNA/DNA-binding. Biochem. Biophys. Res. Commun. 460, 281–286 (2015).

35. Hase, K. et al. RNautophagy/DNautophagy possesses selectivity for RNA/DNA substrates. Nucleic Acids Res. 43, 6439–6449 (2015).

36. Hu, B. et al. Binding of the pathogen receptor HSP90AA1 to avibirnavirus VP2 induces autophagy by inactivating the AKT-MTOR pathway. Autophagy 11, 503–515 (2015).

37. Bryk, M., Banerjee, M., Conte, D.,Jr & Curcio, M. J. The Sgs1 helicase of Saccharomyces cerevisiae inhibits retrotransposition of Ty1 multimeric arrays. Mol. Cell. Biol. 21, 5374–5388 (2001).

38. Ott, K. M., Nguyen, T. & Navarro, C. The DExH box helicase domain of spindle-E is necessary for retrotransposon silencing and axial patterning during Drosophila oogenesis. G3 (Bethesda) 4, 2247–2257 (2014).

39. Taylor, M. S. et al. Affinity proteomics reveals human host factors implicated in discrete stages of LINE-1 retrotransposition. Cell 155, 1034–1048 (2013).

40. Wu-Scharf, D., Jeong, B., Zhang, C. & Cerutti, H. Transgene and transposon silencing in Chlamydomonas reinhardtii by a DEAH-box RNA helicase. Science 290, 1159–1162 (2000).

41. Ranoa, D. R. et al. Cancer therapies activate RIG-I-like receptor pathway through endogenous non-coding RNAs. Oncotarget 7, 26496–26515 (2016).

42. Yun, S. et al. Targeting epigenetic pathways in acute myeloid leukemia and myelodysplastic syndrome: a systematic review of hypomethylating agents trials. Clin. Epigenetics 8, 68-016-0233-2. eCollection 2016 (2016).

43. Jan, M. et al. Clonal evolution of preleukemic hematopoietic stem cells precedes human acute myeloid leukemia. Sci. Transl. Med. 4, 149ra118 (2012).

44. Corces-Zimmerman, M. R., Hong, W. J., Weissman, I. L., Medeiros, B. C. & Majeti, R. Preleukemic mutations in human acute myeloid leukemia affect epigenetic regulators and persist in remission. Proc. Natl. Acad. Sci. U. S. A. 111, 2548–2553 (2014).

45. Corces-Zimmerman, M. R. & Majeti, R. Pre-leukemic evolution of hematopoietic stem cells: the importance of early mutations in leukemogenesis. Leukemia 28, 2276–2282 (2014).

46. Haas, K. et al. Expression and prognostic significance of different mRNA 5'-end variants of the oncogene EVI1 in 266 patients with de novo AML: EVI1 and MDS1/EVI1 overexpression both predict short remission duration. Genes Chromosomes Cancer 47, 288–298 (2008).

47. Yamamoto, H. et al. miR-133 regulates Evi1 expression in AML cells as a potential therapeutic target. Sci. Rep. 6, 19204 (2016).

48. Talpaz, M., Hehlmann, R., Quintas-Cardama, A., Mercer, J. & Cortes, J. Re-emergence of interferon-alpha in the treatment of chronic myeloid leukemia. Leukemia 27, 803–812 (2013).

49. Ortiz, A. & Fuchs, S. Y. Anti-metastatic functions of type 1 interferons: Foundation for the adjuvant therapy of cancer. Cytokine 89, 4–11 (2017).

50. Hasselbalch, H. C. A new era for IFN-alpha in the treatment of Philadelphia-negative chronic myeloproliferative neoplasms. Expert Rev. Hematol. 4, 637–655 (2011).

